# Exploring the structure and mechanics of thin supported inverse bicontinuous cubic phase lipid films

**DOI:** 10.1101/2021.03.29.437497

**Authors:** Andrea Ridolfi, Ben Humphreys, Lucrezia Caselli, Costanza Montis, Tommy Nylander, Debora Berti, Marco Brucale, Francesco Valle

## Abstract

Inverse bicontinuous cubic phase membranes are ubiquitous in nature but their properties and functions are still not fully understood. To shed light on this topic, we herein realize thin supported cubic phase lipid films, characterize their structure and provide the first study of the mechanical properties of these non-lamellar architectures.

---

Lipid self-assembly encompasses a plethora of different structures; inverse bicontinuous cubic (Q_II_) phases are among the most intriguing and at the same time less understood mesophases^1,2^. These structures are characterized by a continuous lipid bilayer membrane that subdivides the three-dimensional space into two interwoven water channels^3^. Lipid Q_II_ phases are ubiquitous in nature^4^, occur spontaneously in stressed or virally infected cells and are involved in numerous biological processes like cell fusion and food digestion^5^. Three types of lipid Q_II_ phase exist: Q_II_ ^P^, Q_II_ ^D^ and Q_II_ ^G^, respectively corresponding to the primitive (P), double diamond (D) and gyroid (G) triply periodical minimal surfaces (TPMS). Compared to planar membranes, the intricate geometry of these architectures provides higher membrane surface-to-volume ratios^6^ which, together with the amphiphilic nature of their lipid components, can be exploited for the encapsulation of hydrophobic, hydrophilic and bioactive molecules, proteins and nanoparticles^7,8^. Thanks to these features, a considerable amount of effort has been put into the study and development of cubic phase nanoparticles, i.e. cubosomes, mostly for drug delivery purposes^9–11^. On the other hand, fewer works in literature focused on the development of homogeneous supported lipid Q_II_ phase films^12–15^. Similarly to Supported Lipid Bilayers (SLBs)^16^, these membrane models could be exploited for characterizing the physicochemical and mechanical properties of natural lipid Q_II_ membranes and applied as biosensing platforms in interface interaction studies.

While the mechanical properties of these nanoarchitectures are still not fully understood, TPMS-inspired macroscopic structures are gaining increasing attention in engineering applications, where the combination of light weight and high impact-and stress-resistance represents a promising solution for multiple structural problems^17–19^. Even if these macroscopic structures and the lipid Q_II_ membranes belong to completely different length scales, they still share the same mathematical description; meaning that perhaps, some of the mechanical properties studied on macroscopic Q_II_ structures might be translated at the nanoscale level, in the study of bioinspired nanomaterials.

In this framework, we herein report one of the first structural characterization of glycerol-monooleate (GMO)-based supported lipid Q_II_^D^ phase films. GMO is known to self-assemble into Q_II_ phases at standard temperature and pressure conditions; moreover, being approved by multiple Health agencies, like the United States Food and Drug Administration and the European Chemicals Agency, it is often used in studies involving non-lamellar membranes^20^. When characterizing the structure of non-lamellar lipid membranes, Small Angle X-ray Scattering (SAXS) is one of the most widely used techniques, since it allows probing large sample areas with high accuracy^21,22^. However, SAXS measurements require specialized instrumentation and expertise (especially when performed at large scale facilities) which make the characterization of these structures a challenging task. In the present study, we also show that it is possible to obtain a comprehensive structural and mechanical characterization of lipid Q_II_^D^ phase films with nanometric thickness, by combining Ellipsometry with Atomic Force Microscopy (AFM). Moreover, by exploiting AFM-based Force Spectroscopy (AFM-FS) we provide the first mechanical characterization of these cubic architectures at the nanoscale level and obtain structural information that are in good agreement with more traditional SAXS measurements performed on the same samples. Compared to classical rheological studies^23–25^, which can only probe the mechanical properties of Q_II_ phase bulk solutions, our analysis describes for the first time the response of Q_II_^D^ membranes to the local deformation introduced by a nanosized object like the AFM tip, closely resembling the mechanical stresses occurring in most biological interactions. In order to obtain nanometric lipid Q_II_^D^ phase films, GMO was dissolved in chloroform, reaching a final concentration of 10 mg/ml and subsequently spin-coated onto the experimental substrate. For SAXS, the GMO solution was directly spin-coated on the Kapton windows of the measuring cells; films were subsequently hydrated and probed using X-rays (see SI for more details). Figure 1 highlights the challenges related to characterizing GMO-based lipid thin films with standard SAXS instrumentation. Traces corresponding to increasingly more concentrated GMO solutions (thus yielding thicker films by spin-coating) progressively show more definite structural peaks. The position of the single peak in the 100 mg/ml sample, corresponds to a lattice parameter of 90 Å while the lattice parameters calculated for the 1000 mg/ml sample and bulk phase are 92.6 ± 0.2 Å and 92.8 ± 0.3 Å, respectively. These results are compatible with the formation of an ordered Q_II_^D^ architecture that is mostly independent of the film thickness; possibly suggesting that the same lipid arrangement is also present in the 10 mg/ml sample, despite it not being detectable in our SAXS setup.

**Figure 1:**
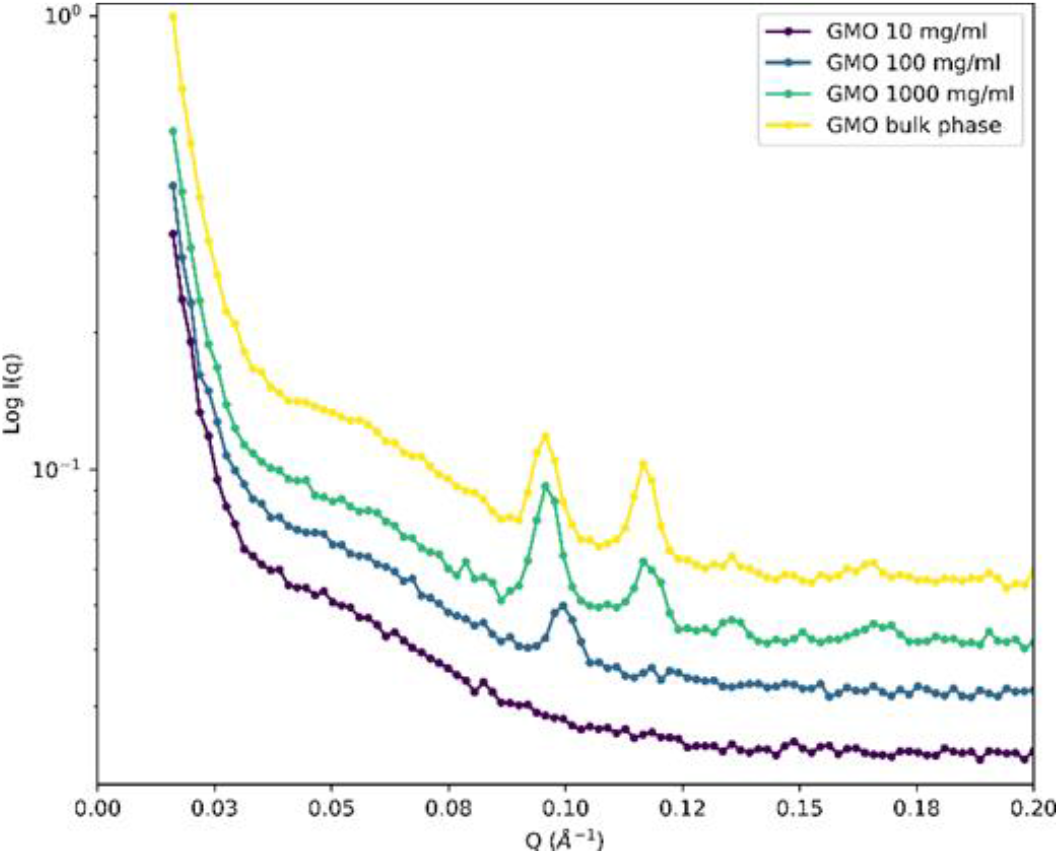
SAXS characterization of lipid Q_II_^D^ phase films at different GMO concentrations. While it is not possible to identify any Bragg peak in the more diluted sample (purple curve), they become more pronounced as the concentration of the sample is increased (until reaching the bulk phase). Peak positions describe the formation of an ordered Q_II_^D^ phase and seem to be independent of the concentration.

Ellipsometry was then used for measuring the thickness and the stability of the 10mg/ml lipid film over time. Since Kapton cannot be probed by polarized visible light, the lipid solution was spin-coated on a silicon substrate, coated with a thin polystyrene layer, which has a surface energy similar to the Kapton substrate. Ellipsometry results reveal the formation of a continuous lipid film presenting a thickness that ranges from 130 to 170 nm which was stable for several hours (please refer to SI) upon hydration.

Even if the low thickness of the 10 mg/ml film hindered its structural characterization by means of SAXS, the cubic architecture can still be probed via AFM imaging; this has already been done by Rittman et al.^13^, who obtained an estimation of the lattice parameter for very similar cubic phase lipid films, spin-coated on Highly Ordered Pyrolytic Graphite (HOPG). Following this approach (see SI for details), we used AFM imaging for obtaining a surface characterization of the lipid film. Image analysis revealed that the probed lipid film presents a cubic architecture, directly exposed to the water interface (Figure 2a). After performing routine image postprocessing procedures on Figure 2a, the calculation of the 2D autocorrelation function (ACF)^26^ (Figure 2b) allowed reducing the noise and estimating a lattice parameter of 97.6 ± 0.34 Å, which is in good agreement with the SAXS analysis and compatible with a Q_II_^D^ phase. The overall morphological properties of the film were assessed via AFM imaging, performed on 10×10 µm^2^ regions (Figure 2c). The film thickness, estimated from the AFM images, is ∼ 150 nm (Figure 2d), in agreement with ellipsometry results.

**Figure 2:**
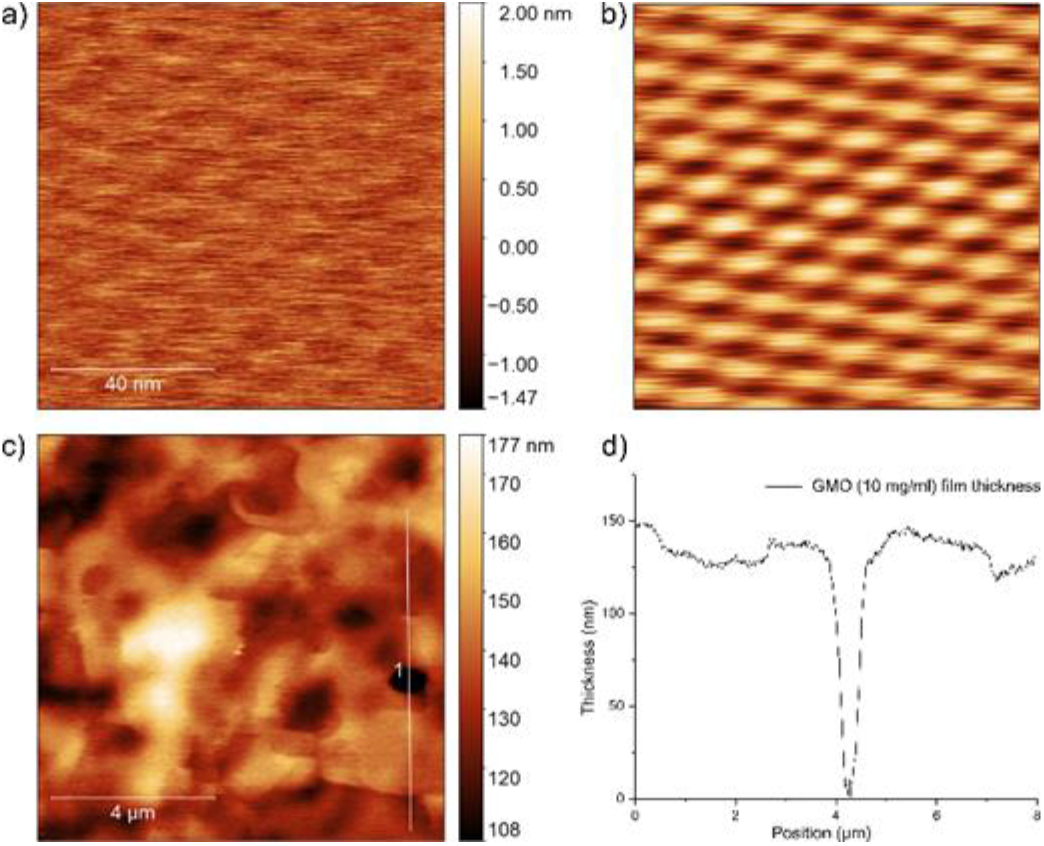
AFM imaging characterization of nanometric GMO-based lipid films: a) the topography of the film surface shows that the film possesses a cubic architecture, exposed to the water interface; b) applying a 2D ACF to the raw AFM image allows denoising and calculating the lattice parameter; c) 10×10 µm2 topography of the lipid film, confirming the presence of a continuous film; d) cut performed on the 10×10 µm2 image allows estimating the thickness of the lipid film, which is approximately 150 nm.

As for SLBs, the herein presented model lipid platforms can be used for studying biological interactions in the presence of highly curved interfaces and for shedding light on the mechanics of these architectures. As a proof of concept, we employed AFM-FS for probing the mechanical properties of the Q_II_^D^ lipid architecture at the nanoscale; an unaddressed topic in the literature regarding non-lamellar lipid membranes. More precisely, AFM-FS was used to investigate the mechanical response of the cubic architecture to a localized indenting force (Figure 3), similar to the ones involved in most biological interactions.

**Figure 3:**
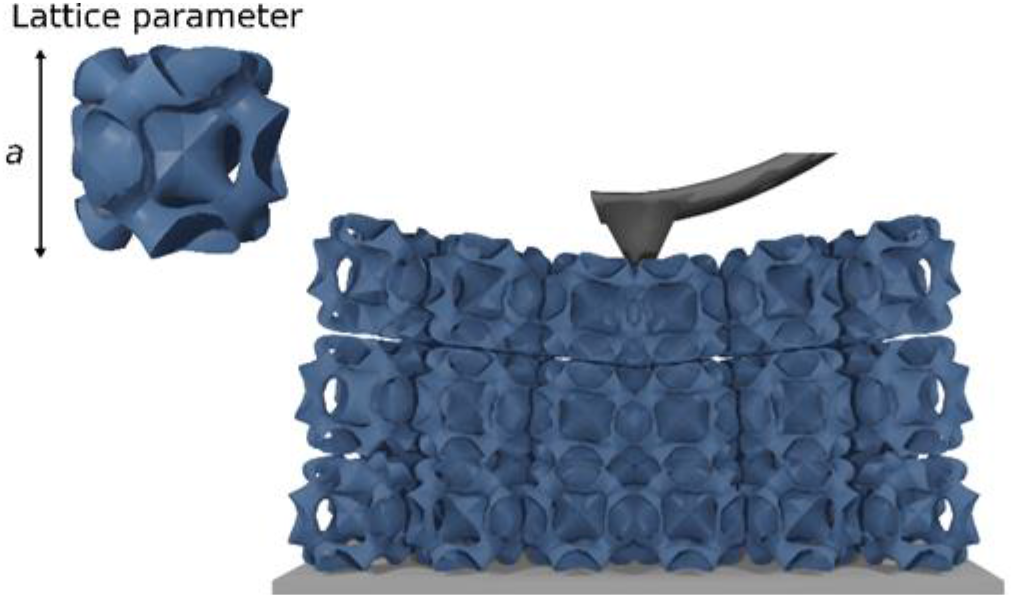
Representation of a typical AFM-FS experiment on a Q_II_^D^ phase film; the AFM tip indents the sample causing a deflection of the cantilever. Recording the forces experienced during the indentation allows determining the mechanical properties of the sample.

In a typical AFM-FS experiment, the forces experienced by the tip while indenting the sample are recorded as a function of its penetration depth and plotted in the so-called force-distance curves (Figure 4a). The mechanical response of the Q_II_^D^ cubic architecture is completely different from that of a classical SLB; after an initial roughly linear regime, all the curves were characterized by a sequence of indentation peaks, describing the sequential mechanical failure of each cubic unit cell.

**Figure 4:**
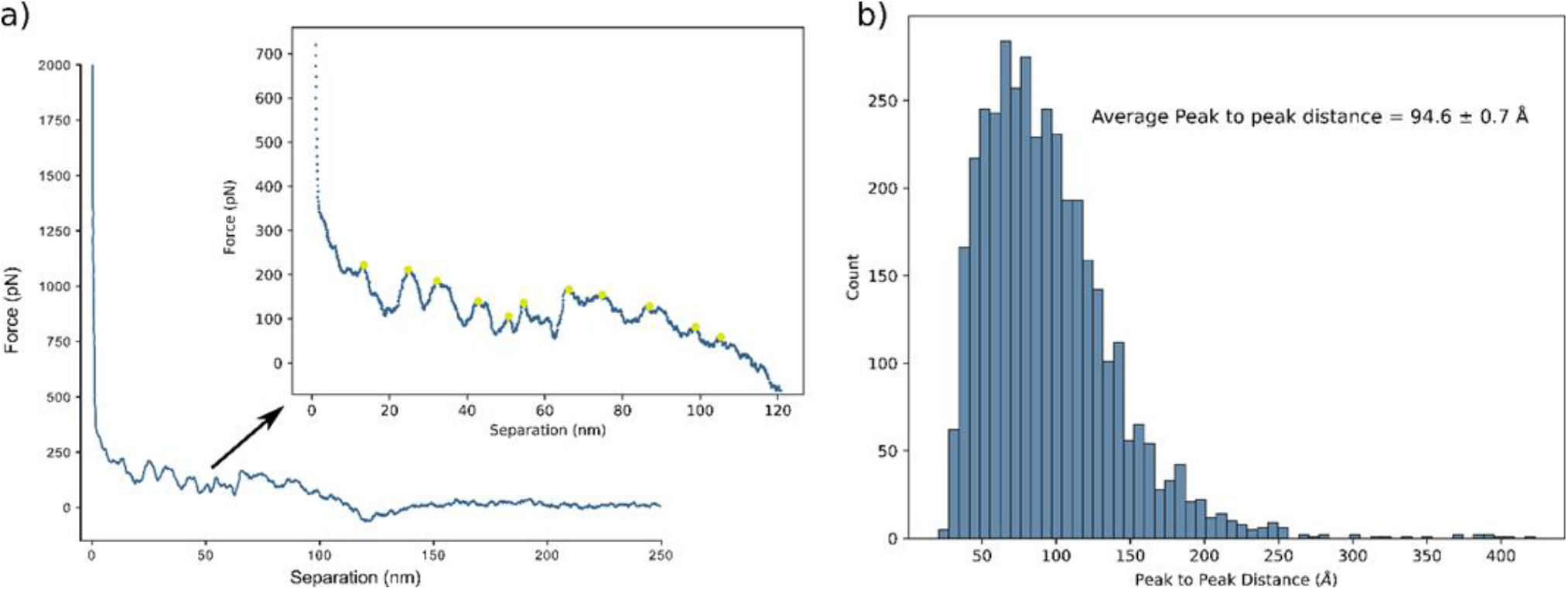
AFM-FS analysis of the lipid Q_II_^D^phase film: a) representative force-distance curve, describing the forces experienced by the AFM tip during the indentation of the cubic architecture; the inset displays the contact portion of the curve with the indentation peaks (green dots) representing the penetration of each cubit unit cell; b) distribution of all the Peak-to-Peak (PtP) distances measured for each pair of peaks, in all the recorded curves; the average PtP distance is 94.6 ± 0.7 Å, in good agreement with the values of lattice parameter obtained from the SAXS and AFM imaging analyses.

Interestingly, this peculiar behavior has been also recorded in multiple mechanical studies on macroscopic TPMS-inspired structures^17–19^, where the stress-strain plots obtained from compression tests are analogous to the inset of Figure 4a, which is reporting the contact part of the whole force-distance curve. This means that these macroscopic structures and the herein probed lipid Q_II_^D^ architecture display a strikingly similar mechanical response to indentation, i.e. a first linear deformation followed by the sequential rupture of each unit cell. It is thus possible that the mechanics of these engineered macroscopic structures is mostly conserved even at the nanoscale. Moreover, the force required to penetrate each unit cell seems to be independent of the penetration depth, meaning that the mechanical resistance of each cubic unit is relatively unaffected by neighbouring ones.

The indentation peaks located along the force-distance curves can also be used for obtaining an alternative characterization of the Q_II_^D^ structure. Indeed, the height of each cubic unit cell also represents the characteristic length scale of the whole architecture, i.e. its lattice parameter (Figure 3). In order to calculate this structural parameter from the force-distance curves, we measured the Peak-to-Peak (PtP) distance for each pair of peaks in all the recorded curves (obtaining a total of 4135 PtP distance values) and analysed their distribution (Figure 4b). The average PtP distance is 94.6 ± 0.7 Å, in good agreement with the lattice parameters from both the SAXS and the AFM imaging analyses (92.6 Å, 92.6 Å and 97.6 Å, respectively). Our analysis of the PtP distance distribution also provides the first example in the literature on the use of AFM-FS for characterizing the structure of non-lamellar lipid membranes. Despite lacking the accuracy of the more traditional scattering techniques (involving both X-rays and/or neutrons)^12,21,22^ and cryo-electron microscopy (cryo-EM)^5,27^, in this context, AFM-FS represents a more accessible and still reliable solution for obtaining a quick structural analysis of these lipid mesophases.

## Conclusions

In the recent years, lipid Q_II_ phases emerged as promising membrane models for multiple biomedical applications; compared to lamellar membranes, their 3D architecture seems to be better suited for the encapsulation of drugs and biomolecules. For this reason, most of the studies available in literature focused on Q_II_ phase nanoparticles, called cubosomes, and fewer efforts were put into the development of Q_II_ phase model platforms, which could have a huge impact in biosensing applications as well as in fundamental studies. This work reports on the fabrication of supported lipid films with nanometric thickness, presenting a Q_II_^D^ architecture. After probing the structure and the stability of the systems with multiple techniques, AFM-FS was used to obtain the first nanomechanical characterization of these membrane models. The mechanical response of the probed lipid Q_II_^D^ phase films revealed interesting analogies with studies performed on similar macroscopic structures, hence suggesting that the response of these architectures to an applied force is independent of their characteristic length scale. AFM-FS also allowed estimating the lattice parameter of the probed Q_II_^D^ membranes, giving results in good agreement with SAXS and therefore offering an alternative and still reliable approach for characterizing the structure of non-lamellar lipid mesophases.

## Acknowledgements

This research has received funding from the Horizon 2020 Framework Programme under the grants FETOPEN-801367 “evFOUNDRY” and FETPROACT-EIC-05-2019 “Bio-Organic Wetsuits”. We thank the SPM@ISMN research facility for support in the AFM experiments.

